# Interaction of Fungal lipase with potential phytotherapeutics

**DOI:** 10.1101/2022.02.11.480159

**Authors:** Farheen Naz, Imran Khan, Asimul Islam, Luqman A Khan

**Affiliations:** Department of Biosciences, Faculty of Natural Sciences, Jamia Millia Islamia, New Delhi-110025; Deoartment of Computer Unit,Qassim Unversity, Buraidah, 51452, KSA; Centre for Interdisciplinary Research in Basic Sciences, Jamia Millia Islamia, Jamia Nagar, New Delhi 110025, India

## Abstract

Interaction of thymol, carvacrol and linalool with fungal lipase and Human Serum Albumin (HSA) have been investigated employing UV-Vis, Fluorescence and Circular dichroism spectroscopy along with docking studies. Thymol, carvacrol and linalool displayed approximately 50% inhibition at 1.5 mmol/litre concentrations using para-nitrophenyl palmitate. UV-Vis spectroscopy give evidence of the formation of lipase-linalool, lipase-carvacrol and lipase-thymol complex at the ground state. Three molecules also showed complex formation with HSA at the ground state. Fluorescence spectroscopy shows strong binding of lipase to thymol (K_a_ of 2.6 x 10^9^ M^-1^) as compared to carvacrol (4.66 x 10^7^ M^-1^) and linalool (5.3 x 10^3^ M^-1^). Number of binding sites showing stoichiometry of association process on lipase is found to be 2.52 (thymol) compared to 2.04 (carvacrol) and 1.12 (linalool). Secondary structure analysis by CD spectra results, following 24 hours incubation at 25°C, with thymol, carvacrol and linalool revealed decrease in negative ellipticity for lipase indicating loss in helical structure as compared with the native protein. The lowering in negative ellipticity was in the order of thymol > carvacrol > linalool.

Results of Fluorescence and CD spectroscopy taken together suggests that thymol and carvacrol are profound disrupter of lipase structure.

Fluorescence spectra following binding of all three molecules with HSA caused blue shift which suggests the compaction of the HSA structure. Association constant of thymol and HSA is 9.6 x 10^8^ M^-1^ which along with ‘n’ value of 2.41 suggests strong association and stable complex formation, association constant for carvacrol and linalool was in range of 10^7^ and 10^3^ respectively.

Docking results give further insight into strong binding of thymol, carvacrol and linalool with lipase having free energy of binding as -7.1 kcal/mol, -5.0 kcal/mol and -5.2 kcal/mol respectively.

To conclude, fungal lipases can be attractive target for controlling their growth and pathogenicity. Employing UV-Vis, Fluorescence and Circular dichroism spectroscopy we have shown that thymol, carvacrol and linalool strongly bind and disrupt structure of fungal lipase, these three phytochemicals also bind well with HSA. Best anti-lipase molecules based on disruption of lipase structure and HSA structure conservation is thymol.

## 1. Introduction

Fungal infections continue to increase with rise in population with predisposing factors, most frequently documented fungal pathogens are *Candida* and *Aspergillus* spp. Commercially available classes of antifungals like polyenes, azoles, allylamines, echinocandins and nucleotide base analogs are associated with severe toxicity and pathogens are becoming increasingly resistant to them. Adverse side effects of current antifungals have promoted scouting for safe and effective natural molecules which can be developed as antifungals directed against fungal growth or pathogenesis.

Secreted lipases are being increasingly implicated in fungal pathogenesis. They are shown to be responsible for destruction of host epidermal and epithelial tissues lipophilic fungal pathogens heavily rely on secreted lipases for their nutrition and growth (1). Lipases are known to play role in fungal adhesion, colonization and persistence (2). Fungal lipases are thus an attractive target for controlling its growth and pathogenesis.

Several phytochemicals are reported to affect fungal growth pathogenesis, reported mechanism of some of them may also involve lipase activity. In this study we select three phytochemicals thymol, carvacrol and linalool which affect fungal virulence or growth and check their direct inhibiting action on *Aspergillus niger* lipase at various concentration.

Linalool (Fig 1 A) exert, antifungal activity by disrupting membrane integrity and arresting the cell cycle of planktonic *C. albicans* (3). In addition, it inhibits *C. albicans* germ tube formation and biofilm formation (4). Thymol (Fig 1 B) appears to bind to the ergosterol in the membrane, which increases ion permeability and ultimately results in cell death (5). Other action mechanisms may be involved, such as the inhibition of spore germination, fungal proliferation and cell respiration (6). Carvacrol (Fig 1 C) acts by interacting with the fungal cell membrane sterols, making it permeable (7).

**Fig 1 A.**
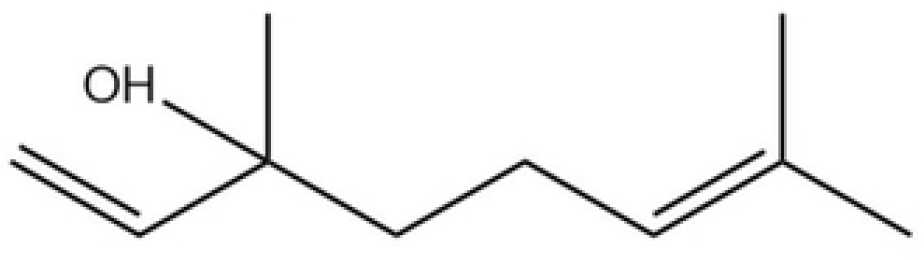
(linalool)

**Fig 1 B.**
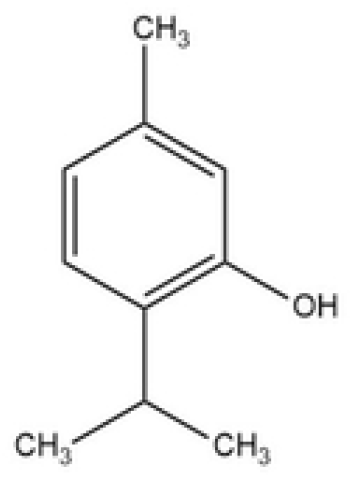
(thymol)

**Fig 1 C.**
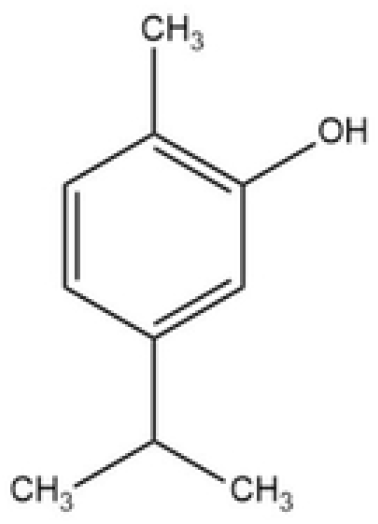
(carvacrol)

For a molecule to be employed as a therapeutic it is required to be a profound disrupter of target protein structure. In addition, it must also be a carrier proteins like Human Serum Albumin (HSA) (Fig 2) but not alter their structure at concentration of interest. Based on above premise, we have investigated interaction of thymol, carvacrol and linalool with Aspergillus niger lipase and HSA employing UV-Visible, Fluorescence and Circular dichroism spectroscopy along with Molecular docking studies. Results obtained have been analyzed for identifying best potential therapeutic molecule out of three lipase inhibitors.

**Fig 2.**
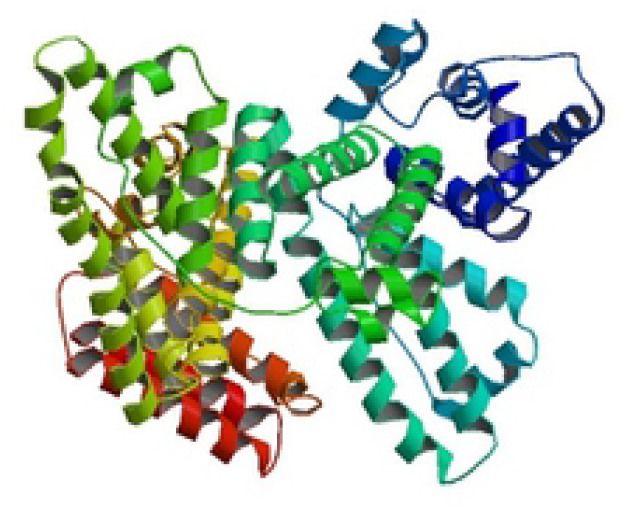
Structure of HSA.

## 2. Materials and method

### 2.1 Materials

Carvacrol, linalool, thymol, *Aspergillus niger* lipase (lipase), Human serum albumin (HSA) and p-nitrophenyl palmitate (pNPP) were purchased from Sigma-Aldrich Chemicals company (USA), gum Arabic from Fisher Scientific, sodium taurocholate from Himedia. All other reagents were of analytical grade and obtained from Merck (India), doubly distilled water was used in all experiments.

### 2.2. Methods

#### 2.2.1 Preparation of solutions

Lipase and HSA were dialyzed against KCl-buffer (pH 7) for 24 hrs. Stock solution of lipase was prepared in 10 mM phosphate buffer at pH 7.4 to obtain a particular concentration. Carvacrol, linalool, thymol was dissolved in small volume of DMSO and stock solution of 1mg/ml was made hrs before use in 10mM phosphate buffer at pH 7.4. Samples were made by mixing the individual protein with increasing volume of carvacrol, linalool, thymol respectively. These samples were incubated at 298.15 K for about 24 hrs.

#### 2.2.2 Lipase inhibition assay

Lipase assays were performed in a 96-well, clear, flat-bottomed plate with 200 μl reaction volume. pNPP was used as a substrate with a reaction buffer of 50 mM sodium phosphate, 5 mM sodium taurocholate, and 10% isopropanol at pH 8.0 (8, 9). Lipase assays used a 200 μl reaction volume and substrate conversion was monitored with a Multiskan Go spectrophotometer (Thermo Scientific) at 410 nm. All assays were run at 37°C and reported results are the average of three replicates that were blank subtracted. The Lipase inhibitory activity was expressed as percentage inhibition calculated as: % Inhibition = [(A_control_ – A_test_ sample)/A_ref_]*100.

#### 2.2.3 Fluorescence measurement

Fluorescence spectra were recorded on Hitachi F-4010 spectrofluorometer equipped with circulating water bath to maintain the temperature of the cell. The quartz cell of path length 1.0 cm and band slit of 5.0 nm were used for all experiments. The excitation wavelength was 280 nm for lipase and HSA. Emission spectra were recorded from 300 – 450 nm for lipase and HSA protein.

Herein, the fluorescence intensities were corrected for absorption of the exciting light and reabsorption of the emitted light to decrease the inner filter effect according to the following equation(10) (Chi & Liu, 2011):

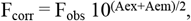

Where F_corr_ and F_obs_ are corrected and experimentally measured fluorescence intensities, respectively. A_ex_ and A_em_ are measured changes in absorbance at λ_ex_ (280 nm) and λ_em_ (343 nm) of linalool, carvacrol and thymol. The fluorescence intensity utilized in this paper was the corrected intensity.

#### 2.2.4 UV–visible absorbance measurement

UV-visible absorbance spectra were obtained using JASCO, V-560 spectrophotometer in the wavelength range 240 to 340 nm. The baseline was corrected for each spectrum.

#### 2.2.5 Circular dichroism (CD) measurement

CD spectra were obtained over wavelength range of 200-250 nm on Jasco-1500 CD Spectrometer equipped with peltier type temperature controller. 0.1cm quartz cell were used to measure far UV CD spectrum. The concentration of protein used was 0.2 mg/ml (lipase) and 0.35 mg/ml (HSA). Each spectrum was scanned thrice and finally average of 3 was used to analyse the results.

#### 2.2.6 Molecular docking

Energy minimized carvacrol, linalool, thymol (ligands) was docked into the active site of lipase using Autodock software (version 4.2). Primarily Autodock (blind docking) required input file in format of pdbqt of ligand and protein. Calculations were carried with the Lamarckian Genetic Algorithm, ligand binding site analysis was visualised using PyMOL (11). The in-silico study provides knowledge about the different binding modes, binding locations and specific residues of the protein involved in interaction with selected molecules. Docking was performed by using AutoDock 4.2 docking software (12) with the help of AutoDock Tools (ADT). The possible binding conformation of the linalool, carvacrol and thymol with proteins was computed by using Lamarckian Genetic Algorithm (LGA) implemented in AutoDock 4.2 (13). *Aspergillus niger* lipase structure was hypothetically demonstrated by I-TASSAR method. The crystal structure of proteins was obtained from RCSB Protein Data Bank with PDB ID; 1aO6 (14) for HSA. The structure of carvacrol, linalool, thymol were sketched on ChemDraw and converted into the 3-D structure by using online SMILES translator. Initially, protein molecule was prepared for docking by removal of water molecules, addition of polar hydrogen atoms and computing Gasteiger charges. Likewise, natural molecule was prepared by calculating their initial positions, orientations, and torsions.

## 3. Results

### 3.1 Effect of linalool, carvacrol and thymol on lipase activity

Fig 3 shows percentage inhibition of lipase activity with increase in linalool, carvacrol and thymol concentration ranging from 0.5 mmoles/litre to 1.5 mmoles/litre using para-nitrophenyl palmitate as substrate. Linalool, carvacrol and thymol markedly inhibited lipase activity. Significant inhibition was obtained at 0.5 mmoles/litre. For all three molecules indicating structural changes begin well below this concentration. According to all binding experiments have been performed in concentration range at 50 μM to 400 μM.

**Fig 3.**
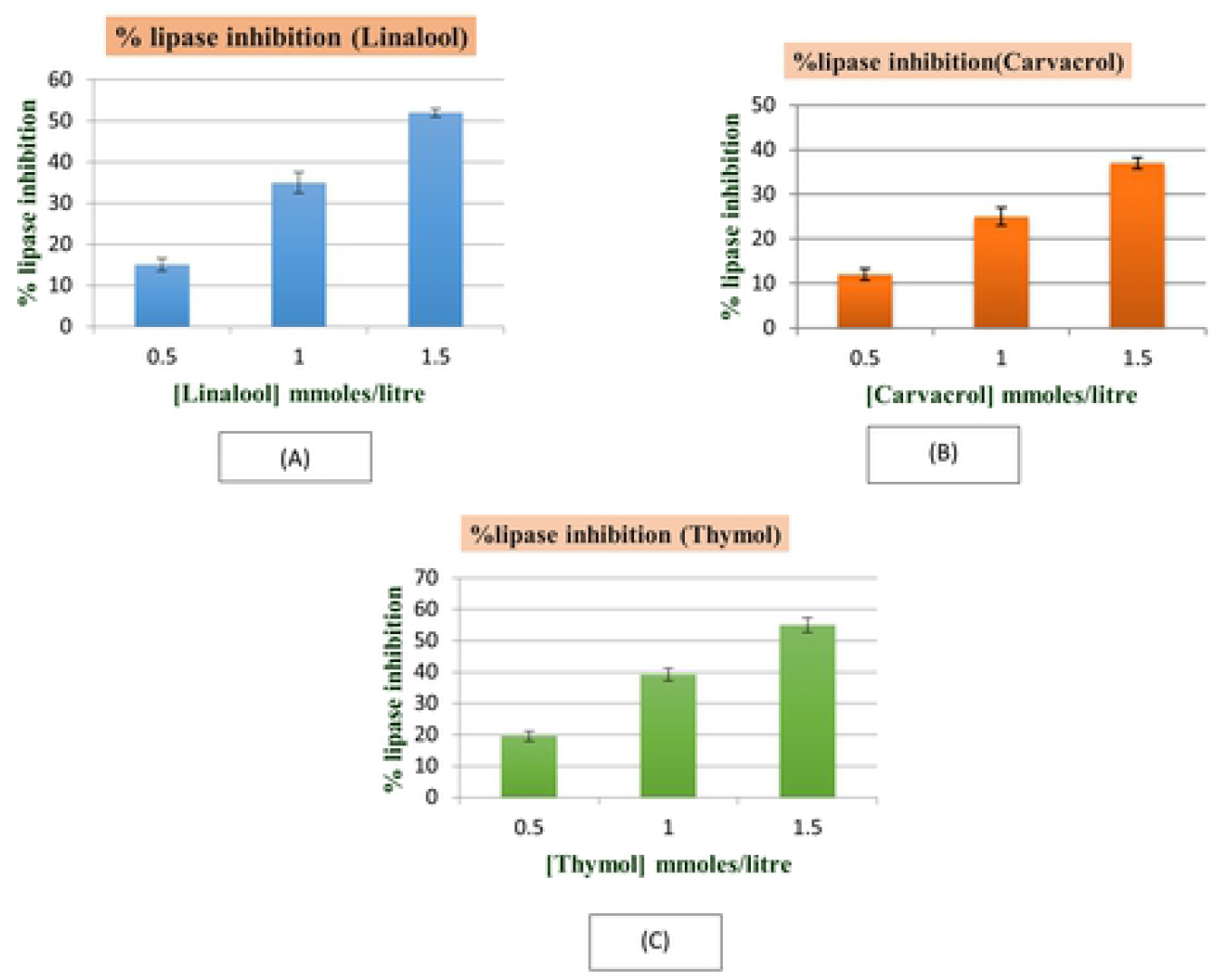
Inhibitory effect of (A) linalool, (B) carvacrol and (C) thymol on lipase activity.

### 2.3 UV–Vis spectroscopy

#### UV–Vis spectroscopy of Lipase with linalool, carvacrol and thymol

UV-Vis absorbance spectra of lipase (0.20 mg/ml) in absence and presence of different concentrations of linalool, carvacrol and thymol were taken from 340 nm to 240 nm (Fig 4A, B and C). Lipase exhibited a strong absorption peak at 278 nm, this band was used for monitoring the interaction between lipase and linalool, carvacrol and thymol. Absorbance of lipase increased regularly with increasing concentration of linalool, carvacrol and thymol indicating formation of ground state lipase-linalool, lipase-carvacrol and lipase - thymol complex. No shift of band is observed from 340 nm to 240 nm suggesting that signal change is due to complex formation leading to changes in microenvironment of chromophore groups (15).

**Fig 4.**
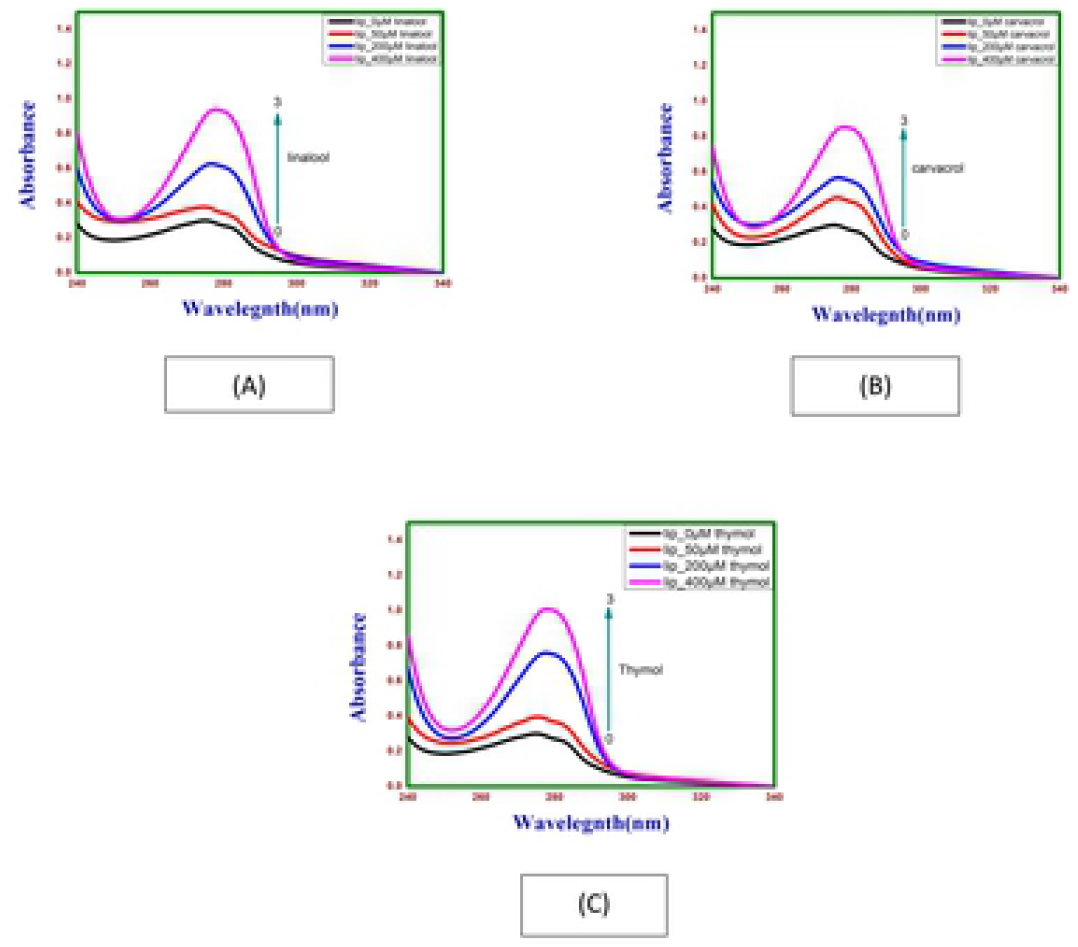
Absorption spectra of lipase as a function of (A) linalool, (B) carvacrol and (C) thymol concentration at 298.15 K and pH 7.4.

### UV–Vis spectroscopy of HSA with linalool, carvacrol and thymol

Absorption of HSA (0.4mg/ml) at different concentrations of linalool, carvacrol and thymol were given as Fig 5(A, B, C) on addition and subsequent increase in the concentration of linalool, carvacrol and thymol absorption peak of HSA showed hyperchromism. This change in the intensity of HSA absorption following addition of linalool, carvacrol and thymol indicate that there are micro-environmental alterations around the protein chromophores upon formation of the HSA-linalool, HSA-carvacrol and HSA-thymol complex. Maximum peak position of linalool–HSA, carvacrol–HSA and thymol–HSA was observed to be shifted slightly towards higher wavelength region from 278 to 278.5, 278.2 and 278.4 nm respectively.

**Fig 5.**
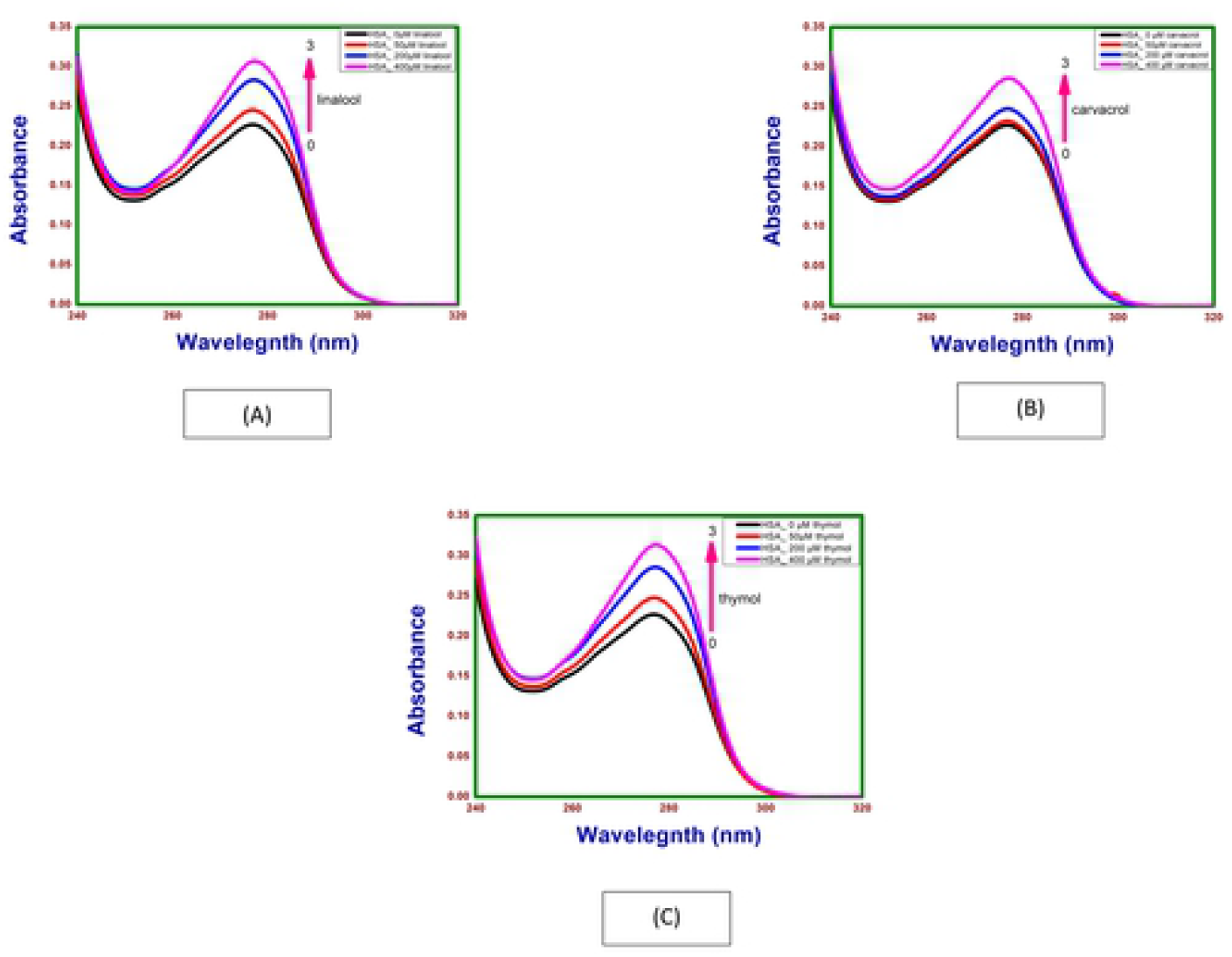
Absorption spectra of HSA as a function of (A) thymol, (B) carvacrol, and (C) linalool concentrations at 298.15 K and pH 7.4.

### 3.3 Fluorescence spectroscopy studies

#### Effect of linalool, carvacrol and thymol on fluorescence spectra of lipase

Fig 6(A, B and C) shows fluorescence emission spectra of lipase in the absence and presence of different concentrations of linalool, carvacrol and thymol (50 μM – 400 μM). A gradual decrease in the fluorescence intensity was observed following addition of small aliquots of linalool, carvacrol and thymol to fixed volume of lipase. Emission fluorescence spectra were measured in the 300-450 nm interval, at a fixed excitation wavelength of 280 nm. Occurrence of an emission maximum at 344 nm in the fluorescence spectrum of lipase can be attributed to the presence of tryptophan residue of lipase. Even though the intrinsic fluorescence of pancreatic lipase is attributed to three amino acid residues, phenylalanine (Phe), tyrosine (Tyr), and tryptophan (Trp), the fluorescence of Phe and Tyr residues at this wavelength is negligible, so intrinsic fluorescence of the fungal enzyme is mainly due to Trp (16). Addition of linalool, carvacrol and thymol to lipase individually produced significant quenching in the Trp fluorescence intensity in a concentration dependent manner without any shift in the emission maximum throughout the titration. Free linalool, carvacrol and thymol did not fluorescence near the emission maximum of lipase. Such decrease in the fluorescence intensity of lipase in the presence of linalool, carvacrol and thymol is suggestive of the interaction of linalool, carvacrol and thymol with lipase. Several earlier reports have shown similar fluorescence quenching of protein upon interaction with various drug molecules, without any shift in the emission maximum (17). The observed quenching in the protein fluorescence can be induced by a variety of molecular interactions such as rearrangement of molecules, ground-state complex formation, excited-state reactions, energy transfer and collisional quenching (18).

**Fig 6.**
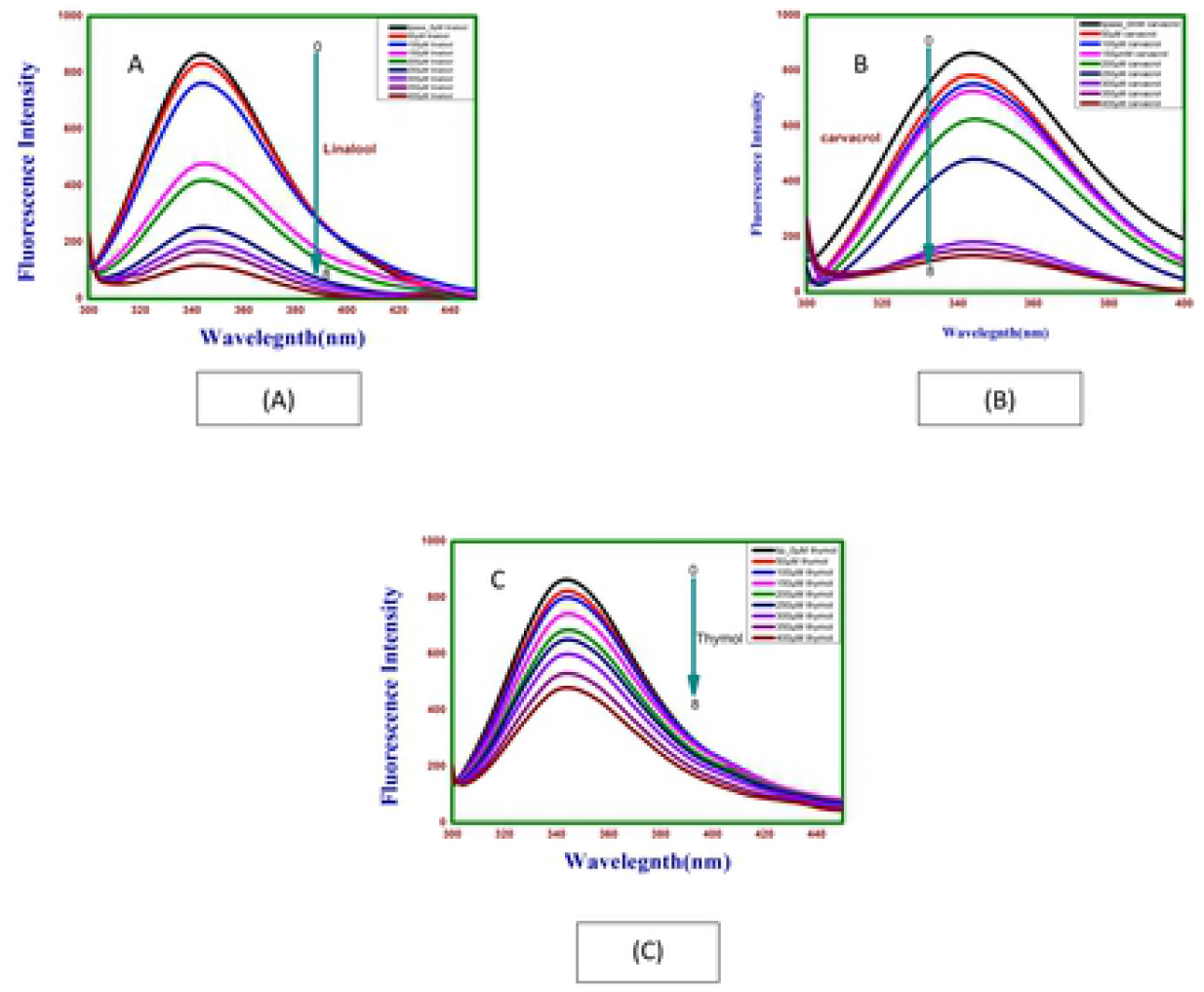
Emission spectra of lipase on (A) linalool, (B) carvacrol and (C) thymol at different concentrations.

*k_sv_* is Stern–Volmer constant; *k_q_* is double molecular dynamic quenching rate constant; R^2^ is correlation coefficient, *K_a_* is binding constant; *n* is stoichiometry of association process

In order to gain insight into quenching mechanism involved in linalool-lipase, carvacrol-lipase and thymol-lipase interaction, or to gain further insight into the mechanism of tertiary structure we analysed the fluorescence quenching data using Stern–Volmer equation.

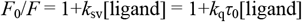

where *F*_0_ and *F* are the fluorescence emission intensities without and with linalool, carvacrol and thymol; [ligand] is ligand concentration in moles/l; *k*_q_ is the fluorescence quenching rate constant; *τ*_0_ is the fluorescence lifetime without quencher and its reference value used is10^-9^s obtained for human pancreatic lipase (19). S1(A-C) Figs. and Table 1. gives *k_sv_* and *k_q_* values of Lipase - lin as 1.73 x 10^4^ M^-1^ and 1.73 x 10^13^ M^−1^s^−1^, Lipase - car as 2.17 x 10^3^ and 2.17 x 10^12^, Lipase - thy as 1.76 x 10^4^ and 1.76 x 10^13^ respectively. Since obtained *k*_q_ value for lipase is greater than the maximum scatter collision quenching constant (2 x 10^10^ M^−1^s^−1^) this indicates static quenching mechanism due to lipase-linalool, lipase - carvacrol and lipase-thymol complex formation rather than dynamic collision(20),. Binding constant and number of binding sites were determined by plotting log (*F*_0_ - *F*)/*F* versus log [ligand] according to equation:

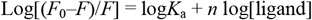

where *K_a_* and *n* are binding constants and stoichiometry of association process respectively. Linear relationship is obtained from log(*F* – *F*_0_) / *F* versus log[ligand]. S2(A-C) Figs. and Table 1 described the values of *K*_a_ of lipase - linalool is equal to 5.3 x 10^3^ M^-1^, lipase - carvacrol to 4.66 x 10^7^ M^-1^ and lipase - thymol 2.6 x 10^9^ M^-1^. Lipase showed strongest affinity for thymol followed by carvacrol and linalool(21). Slope of the plot indicates binding sites on lipase for linalool (*n*=1.12), carvacrol (*n*=2.04) and thymol (*n*=2.52). These ‘n’ values however do not mean number of equivalent binding sites but only stochiometry of association process.

**Table 1:** Stern–Volmer and quenching constants for lipase biomolecular diffusion-controlled quenching constant, binding constant and number of binding sites at pH 7.4.

| Complex | $k_{sv} (\text{M}^{-1})$ | $k_q (\text{M}^{-1}\text{s}^{-1})$ | $R^2$ | $K_a (\text{M}^{-1})$ | $n$ | $R^2$ |
| --- | --- | --- | --- | --- | --- | --- |
| Lipase- lin | $1.73 \times 10^4$ | $1.73 \times 10^{13}$ | 0.96226 | $5.3 \times 10^3$ | 1.12 | 0.99679 |
| Lipase- car | $2.17 \times 10^3$ | $2.17 \times 10^{12}$ | 0.99229 | $4.6 \times 10^7$ | 2.04 | 0.96649 |
| Lipase- thy | $1.76 \times 10^4$ | $1.76 \times 10^{13}$ | 0.94929 | $2.6 \times 10^9$ | 2.52 | 0.99162 |

#### Effect of linalool, carvacrol and thymol on fluorescence spectra of Human Serum Albumin (HSA)

The fluorescence emission spectra of HSA in the absence and the presence of increasing linalool concentrations (50–400 μM with 50 μM intervals) is shown as Fig 7A. The HSA were excited at 280nm and spectra were taken from 300nm to 400nm. Occurrence of an emission maximum of HSA in absence of linalool was at 342 nm which can be attributed to the presence of single tryptophan residue (Trp214) in the subdomain IIA of HSA (22). Addition of linalool to HSA produced significant quenching in the Trp fluorescence emission in a concentration dependent manner with slight blue shift from 342 to 340nm (2nm) in the emission maximum through the titration. Blue shift suggests that the fluorescence chromophore of HSA is placed in a more hydrophobic environment after the addition of linalool which can be generally related to the compaction of the HSA. Similarly, addition of carvacrol and thymol to HSA shows slight blue shift from 342 to 341nm (1nm) and 342 to 340nm (2nm), respectively, in the emission maximum throughout the titration as shown in Fig 7(B and C).

**Fig 7.**
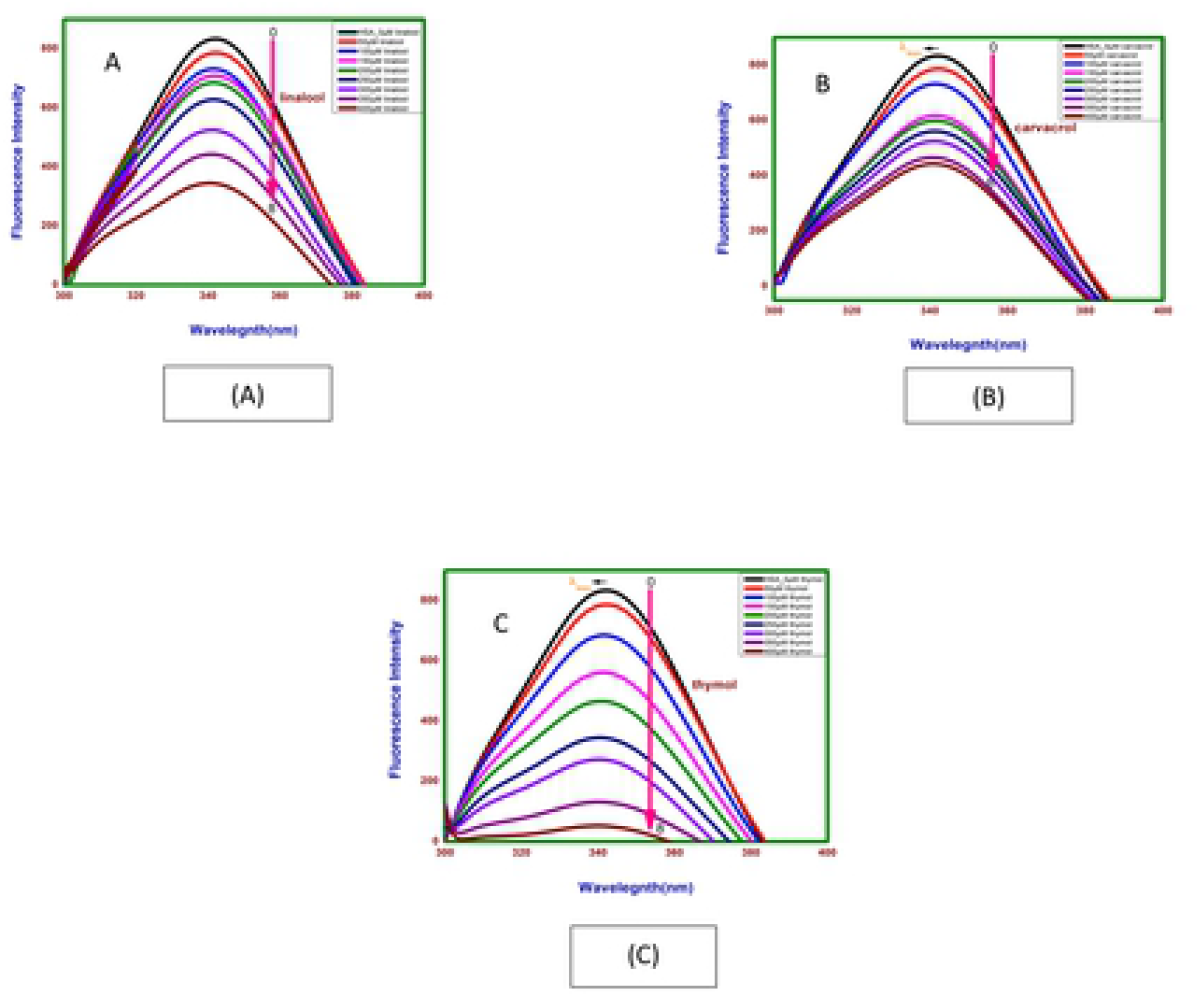
Emission spectra of HSA with (A) linalool, (B) carvacrol and (C) thymol at different concentrations.

The plot of *F*_0_/*F* against [linalool] at 298K is given as S3A Fig. The values of *k_sv_* and *k*_q_ could be calculated from the slope of curves. The average lifetime of the fluorophore in the absence of the ligand as 6.38 × 10^-9^ s for HSA (23). Table 2 and S3A Fig gives the *k*_sv_ and *k*_q_ values of HSA with linalool as3.32 x 10^3^ M^-1^ and 3.32 x 10^12^ M^-1^s^-1^ respectively. Since obtained value of *k*_q_ for HSA is greater than the maximum scatter collision quenching constant i.e. 2 x 10^10^ M^-1^s^-1^, this clearly indicated that this fluorescence quenching mechanism was the static quenching due to linalool - HSA complex formation rather than dynamic collision. *K*_a_ and ‘*n*’ were calculated from equation: Log[(*F*_0_–*F*)/*F*] = log*K*_a_ + *n* log[linalool]. 3.40 x 10^4^ M^-1^ and 1.36 respectively indicating low affinity of linalool for HSA as shown in S4A Fig. and Table 2. Similarly, spectra and plots of Stern-volmer and double logarithm plot for fluorescence intensity following binding of HSA with carvacrol and thymol are given in S3B and C Figs, S4B and C Figs respectively. In both cases the binding was found to static. *k*_sv_ and *k*_q_ determined from stern-volmer equation and *K*_a_ and *n* determined by double - logarithm. Plot for all 3 ligands is summarised as Table 2.

**Table 2.** The value of Stern-Volmer quenching constant (*k*_sv_) and quenching rate constant (*k*_q_), binding constant (*K*_a_) and number of binding site (*n*) in the absence and presence of thymol, carvacrol and linalool following binding to HSA.

| Protein-Molecule | $k_{sv}(M^{-1})$ | $k_q(M^{-1}s^{-1})$ | $R^2$ | $K_a(M^{-1})$ | $n$ | $R^2$ |
| --- | --- | --- | --- | --- | --- | --- |
| HSA-Linalool | $3.32 \times 10^3$ | $3.32 \times 10^{12}$ | 0.96226 | $3.4 \times 10^4$ | 1.36 | 0.99679 |
| HSA-Carvacrol | $2.33 \times 10^3$ | $2.33 \times 10^{12}$ | 0.99229 | $2.2 \times 10^4$ | 1.29 | 0.96649 |
| HSA-Thymol | $3.21 \times 10^4$ | $3.21 \times 10^{13}$ | 0.94929 | $9.6 \times 10^8$ | 2.41 | 0.99162 |

### 3.4 Circular dichroic spectroscopy studies

Far-UV CD spectra has been investigated to characterize the secondary structure and environment around the peptide backbone following binding of linalool, carvacrol and thymol to lipase and HSA. Incubation of linalool, carvacrol and thymol /buffer with proteins in this set of experiments was for 24 hours at 298.15 K. Evaluation of such conformational aspects of drug-protein binding is crucial in assessing the efficacy of the drug as a therapeutic agent.

#### Influence of linalool, carvacrol and thymol binding on secondary structure of lipase

The far-UV CD spectra of lipase in the absence and presence of linalool, carvacrol and thymol (400 μM) were taken from 250 nm to 200 nm (Fig. 8A, B and C and Table 3). In presence of linalool, carvacrol and thymol there was a decrease in MRE values, particularly at 222 nm, which is an index of secondary structure. MRE values decrease from −9.34 to −8.74, −7.71 and −6.69 following binding of linalool, carvacrol and thymol respectively. The lowering in the negative ellipticity hints towards a decrease in the α-helical content and suggests an unfolding of the peptide strand (24).

**Fig 8.**
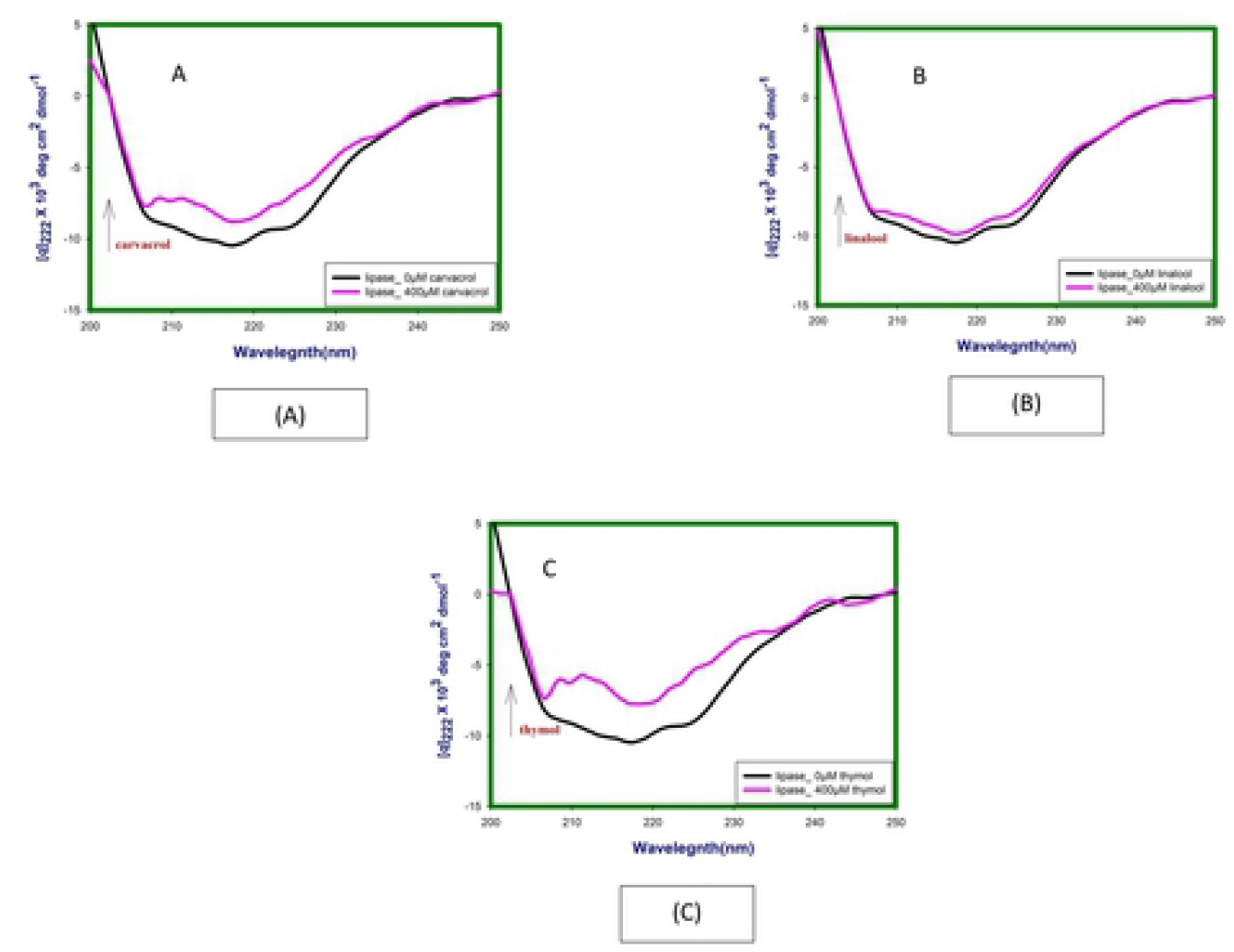
Far-UV CD spectra at 24hrs lipase in absence and presence of (A) linalool, (B) carvacrol and (C) thymol at 298.15 K and pH 7.4.

**Table 3.** summarises MRE values of free and complexed lipase and HSA.

| Enzyme/<br>Protein | MRE x 10<sup>3</sup> at<br>222nm (native) | MRE x 10<sup>3</sup> at 222nm (complex_400 μM) |  |  |
| --- | --- | --- | --- | --- |
|  |  | Thymol | Carvacrol | Linalool |
| Lipase | -9.34 | -6.69 | -7.71 | -8.74 |
| HSA | -17.76 | -23.56 | -24.91 | -23.13 |

#### Influence of linalool, carvacrol and thymol binding on secondary structure of HSA

Far UV CD spectra of HSA in absence and presence of linalool, carvacrol and thymol are given in Table 3 and Fig. 9A, B and C. Upon titration with increasing concentration of linalool, carvacrol and thymol (50 μM, 200 μM and 400 μM) it appears that linalool induces the secondary structure of HSA and increases its helicity. The decreasing in the negative ellipticity from −17.76 (free HSA) to −23.13 (for complex at 400 μM of linalool) at 222nm without any specific shift of the peaks result towards an increase in the α-helical content and suggests folding of the peptide strand.

**Fig 9A.**
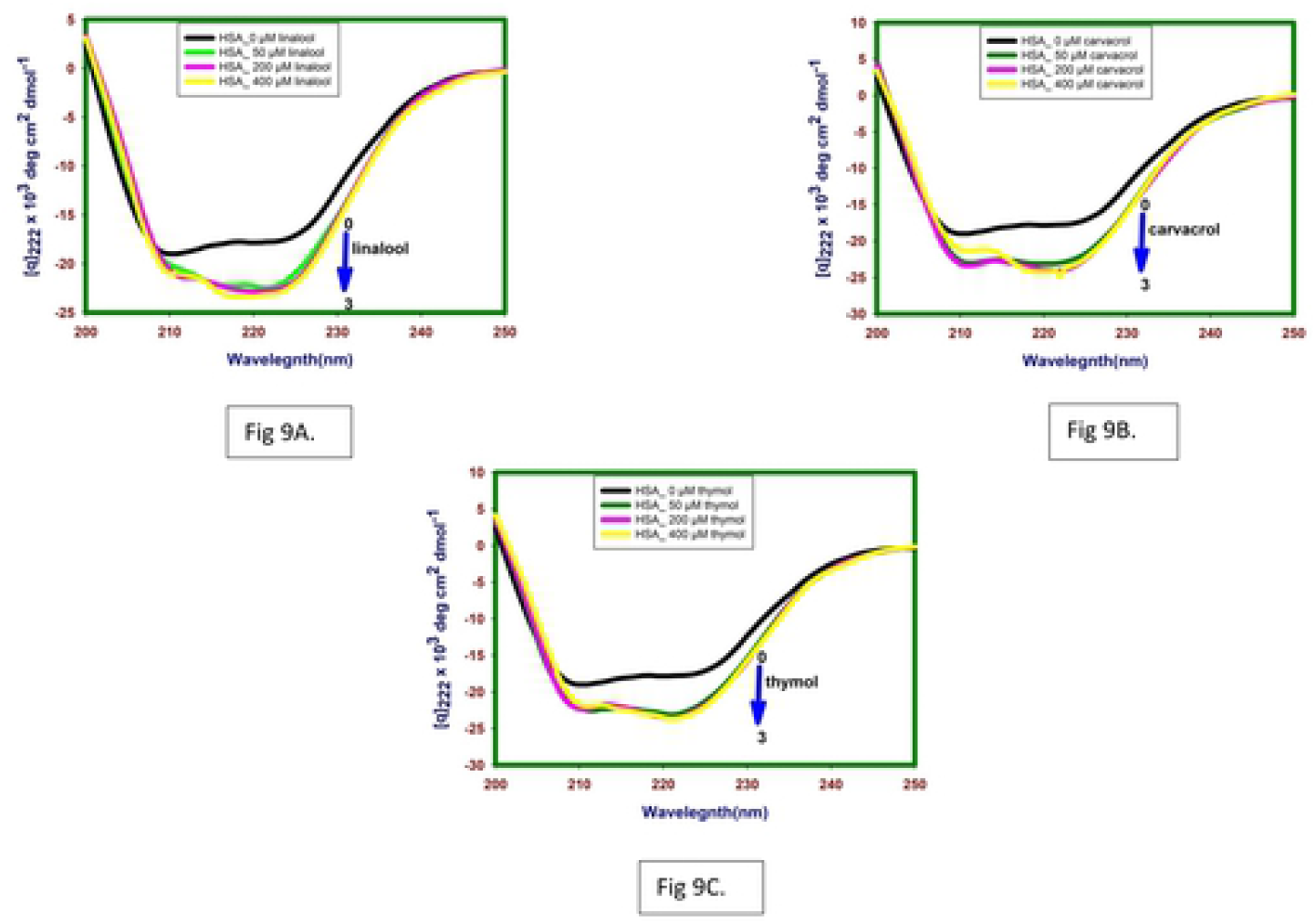
Far-UV CD spectra at 24hrs HSA in absence and presence of linalool, (9B) carvacrol and (9C) thymol at 298.15 K and pH 7.4.

Similar spectra for carvacrol and thymol are given in Fig. 9B and C respectively. The decreasing in the negative ellipticity for carvacrol and thymol −17.76 to −24.91 and −23.56 at 222nm without any specific shift of the peaks suggests increase in the α-helical content and suggests folding of the peptide strand.

### 3.5 Molecular docking results

#### Molecular docking of lipase with linalool, carvacrol and thymol

We use *ab-initio* modeling from I-TASSER (S5 Fig), pair wise sequence alignments were performed using the Needleman-Wunsch algorithm as crystal structure of lipase of *Aspergillus niger* is not available (25). Docking was performed using Pymol program. The result show lowest binding energy of linalool −5.28 kcal/mol for lipase as compared to other two molecules (Table 4). As presented the hydroxyl fragment of the linalool shown to engage the Trp190 via the formation of one polar bond. Whereas, the aromatic moiety of the linalool shown to engage the TRP384, LEU108, TRP110, LEU137, LEU387, ALA138, VAL145 for hydrophobic interaction and for cation-pi interaction Trp190. The other non-bonded interaction was also observed with the involvement of Asp134, Asn380 (Fig 10a and 10b). The similar value of interaction has been also reported on close inspection of the HB plot of the linalool as shown in Fig 10c. HB plot showing hydrogen bonding interaction between lipase and ligand.

**Table 4:**
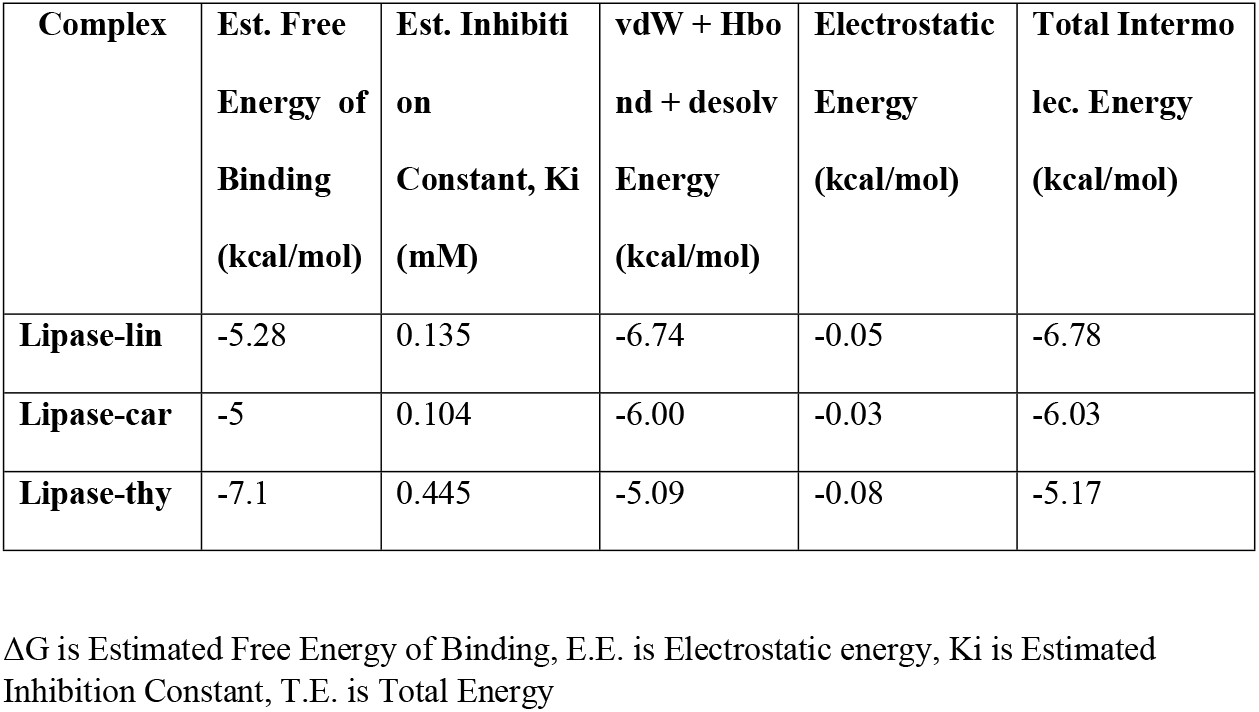
Binding studies of lipase with linalool, carvacrol and thymol by molecular docking.

| Complex | Est. Free Energy of Binding (kcal/mol) | Est. Inhibition Constant, $K_i$ (mM) | vdW + Hbond + desolv Energy (kcal/mol) | Electrostatic Energy (kcal/mol) | Total Inter-molecular Energy (kcal/mol) |
| --- | --- | --- | --- | --- | --- |
| Lipase-lin | -5.28 | 0.135 | -6.74 | -0.05 | -6.78 |
| Lipase-car | -5 | 0.104 | -6.00 | -0.03 | -6.03 |
| Lipase-thy | -7.1 | 0.445 | -5.09 | -0.08 | -5.17 |

$\Delta G$ is Estimated Free Energy of Binding, E.E. is Electrostatic energy, $K_i$ is Estimated Inhibition Constant, T.E. is Total Energy

**Fig 10.**
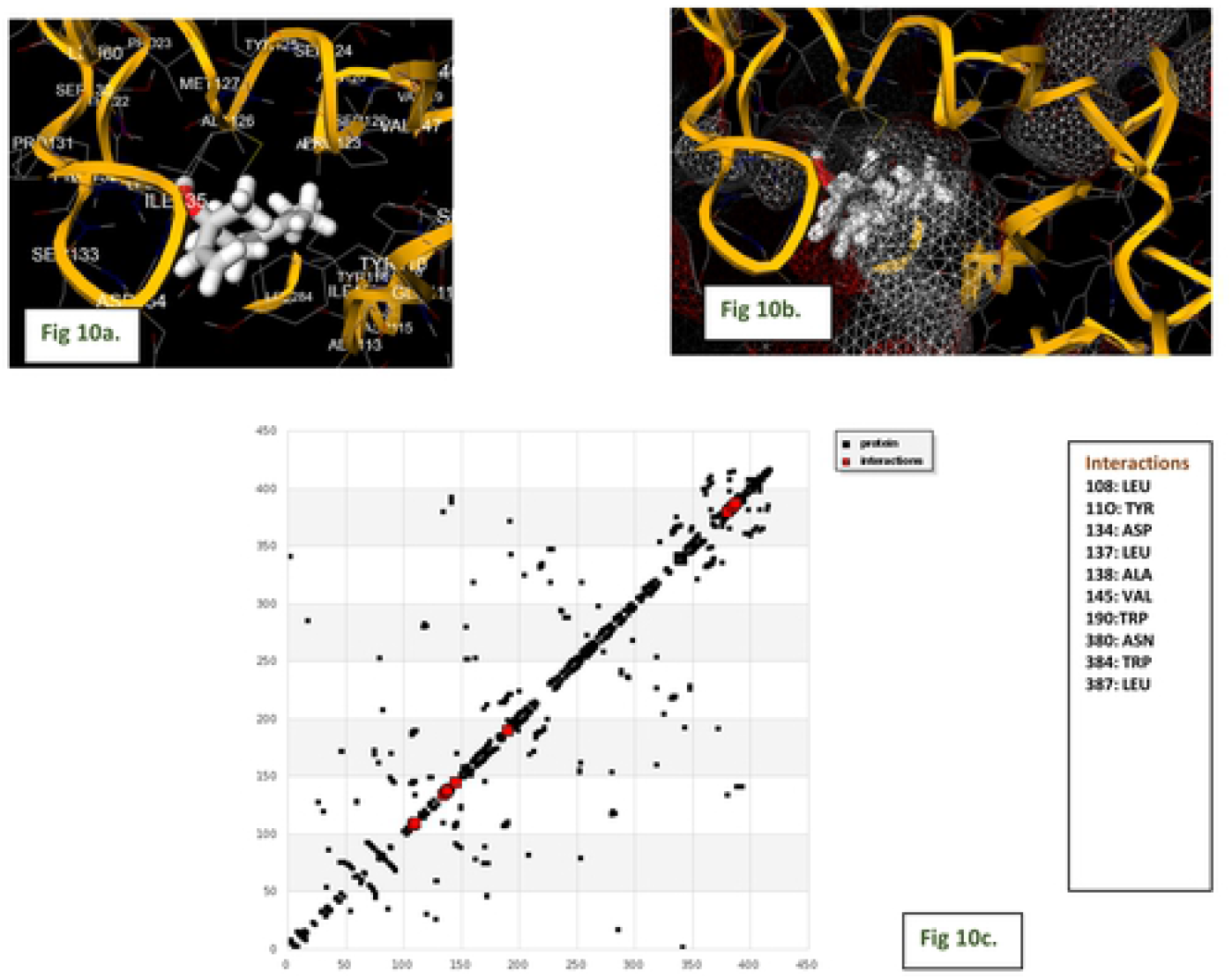
Fig 10(a). Docked orientation of linalool at the binding site of *A niger* lipase with showing bonding residues stick model in yellow and ligand in white color, Fig 10(b). Binding mode of *A niger* lipase with linalool to lipase surface residues network like, Fig 10(c). HB Plot of linalool in the active site of lipase domain.

Carvacrol also makes stable complex with lipase also with binding energy of −5 kcal/mol. It has been found that, due to structural difference in the position of the hydroxyl group in carvacrol, it showed additional bonds with the neighbouring residues, such as, it showed to interact with TRP384, TRP190, ALA138, LEU137, LEU387, LEU108 for hydrophobic interaction and Tyr110, Asp134, Ile135 and Asn380 to create another non bonded interaction (Fig 10 and 10e). The similar value of interaction has been also reported on close inspection of the HB plot of the carvacrol as shown in Fig 10f.

**Fig 10.**
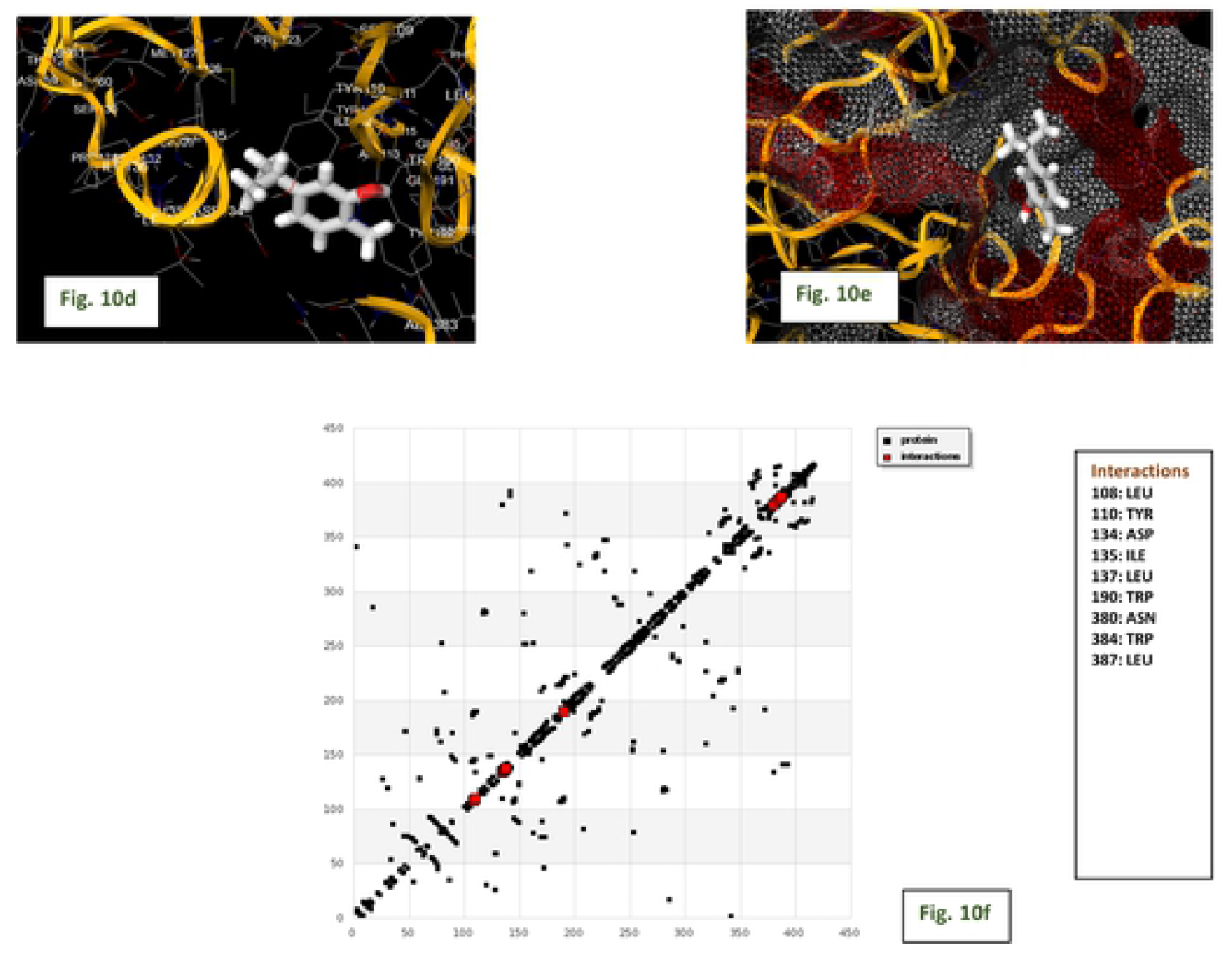
Fig 10(d). Docked orientation of carvacrol at the binding site of *A niger* lipase with showing bonding residues stick model in yellow and ligand in white color, Fig 10(e). Binding mode of *A niger* lipase with carvacrol to lipase surface residues network like, Fig 10(f). HB Plot of carvacrol in the active site of lipase domain.

Interestingly. thymol on docking with lipase shows very prolific engagements with the neighbouring residues as shown in (Fig 10g and 10h). They comparatively give −7.1 kcal/mol of binding energy. The compound was found to be buried in the active site of the binding domain. It shows to engage the Ala113 via the formation of one polar bond. Whereas, the aromatic moiety of the thymol seems to engage the Phe14, Ile112, Ile112, Tyr192 for hydrophobic interaction and for cation-pi interaction Trp110. The other non-bonded interactions were also observed with the involvement of Asp134, Gln376. The similar value of interaction has been also reported on close inspection of the HB plot of the thymol as shown in Fig 10i.

**Fig 10.**
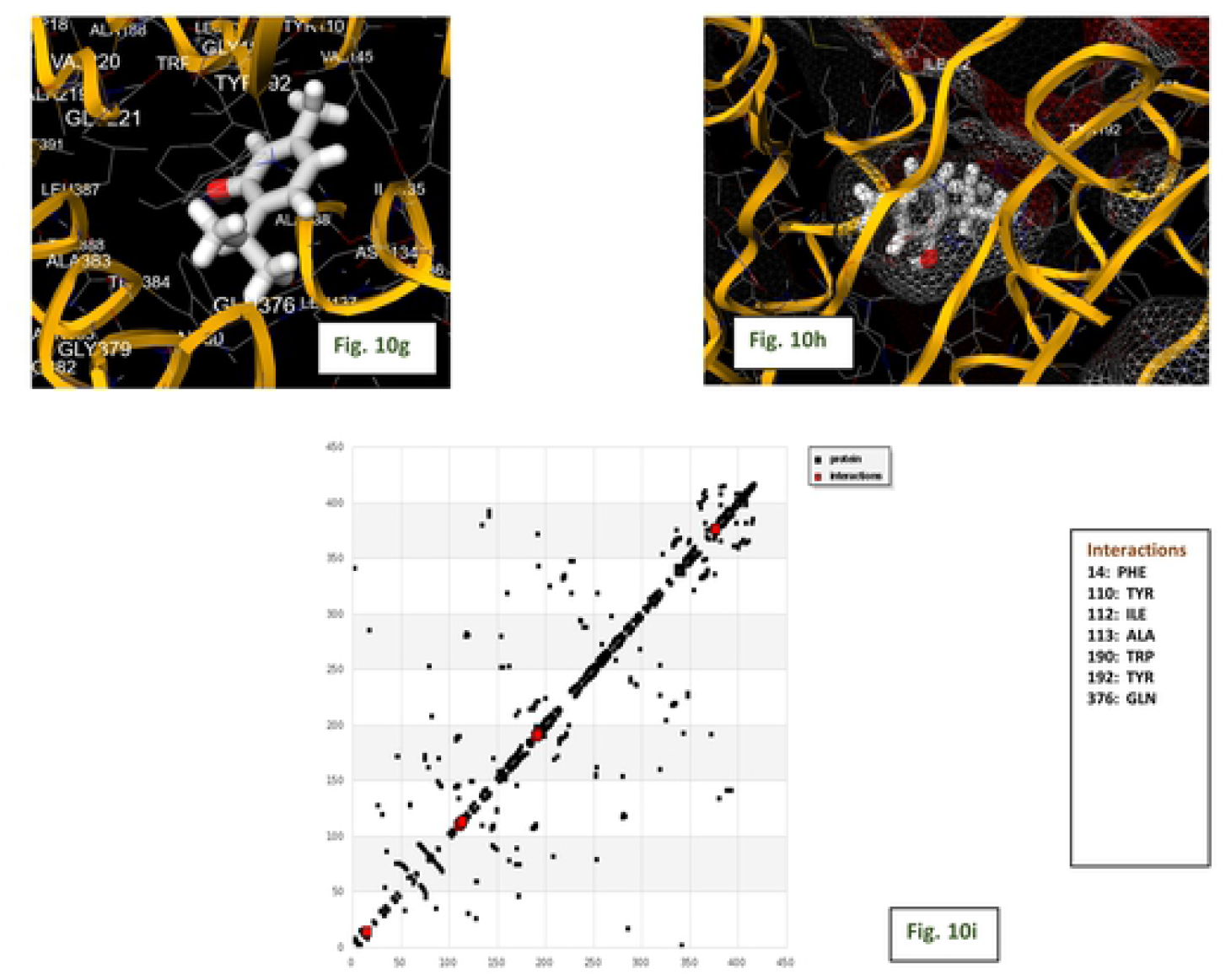
Fig 10(g). Docked orientation of thymol at the binding site of *A niger* lipase with showing bonding residues stick model in yellow and ligand in white color, Fig 10(h). Binding mode of *A niger* lipase with thymol to lipase surface residues network like, Fig 10(i). HB Plot of thymol in the active site of lipase domain.

#### Molecular docking of linalool, carvacrol and thymol with HSA

The binding images of linalool–HSA complex is shown in Fig. 11a and 11b, and the results of free binding energy and other values are given in Table 5. The binding results indicate that linalool is very close to the active site amino acid residues His146, Pro147, Tyr148, Tyr149, Ala194, Arg197, Asp108, Gln459 at site I in subdomain IIA. The hydrogen bonding between linalool and HSA (Tyr149, ARG256) is also responsible for maintaining the stability of this complex. Whereas, the aromatic moiety of the linalool shown to engage the His146, Tyr148, Pro147, Ala194 for hydrophobic interaction and for polar interaction Arg197, Asp108. The other non-bonded interaction was also observed with the involvement of Gln459, Ser193. The similar value of interaction has been also reported on close inspection of the HB plot of the linalool and HSA as shown in Fig 11c. Linalool binds to the binding site of HSA with minimum binding energy (ΔG) −4.21 kcal/mol (Table 5).

**Table 5.** Binding studies of HSA with thymol, carvacrol and linalool by molecular docking.

| Complex | Est. Free Energy of Binding (kcal/mol) | Est. Inhibition Constant, $K_i$ (mM) | vdW + Hbond + desolv Energy (kcal/mol) | Electrostatic Energy (kcal/mol) | Total Inter molec. Energy (kcal/mol) |
| --- | --- | --- | --- | --- | --- |
| HSA-Linalool | -4.21 | 0.818 | -5.68 | -0.11 | -5.79 |
| HSA-Carvacrol | -4.77 | 0.318 | -5.34 | -0.03 | -5.37 |
| HSA-Thymol | -4.77 | 0.318 | -5.29 | -0.1 | -5.41 |

**Fig 11.**
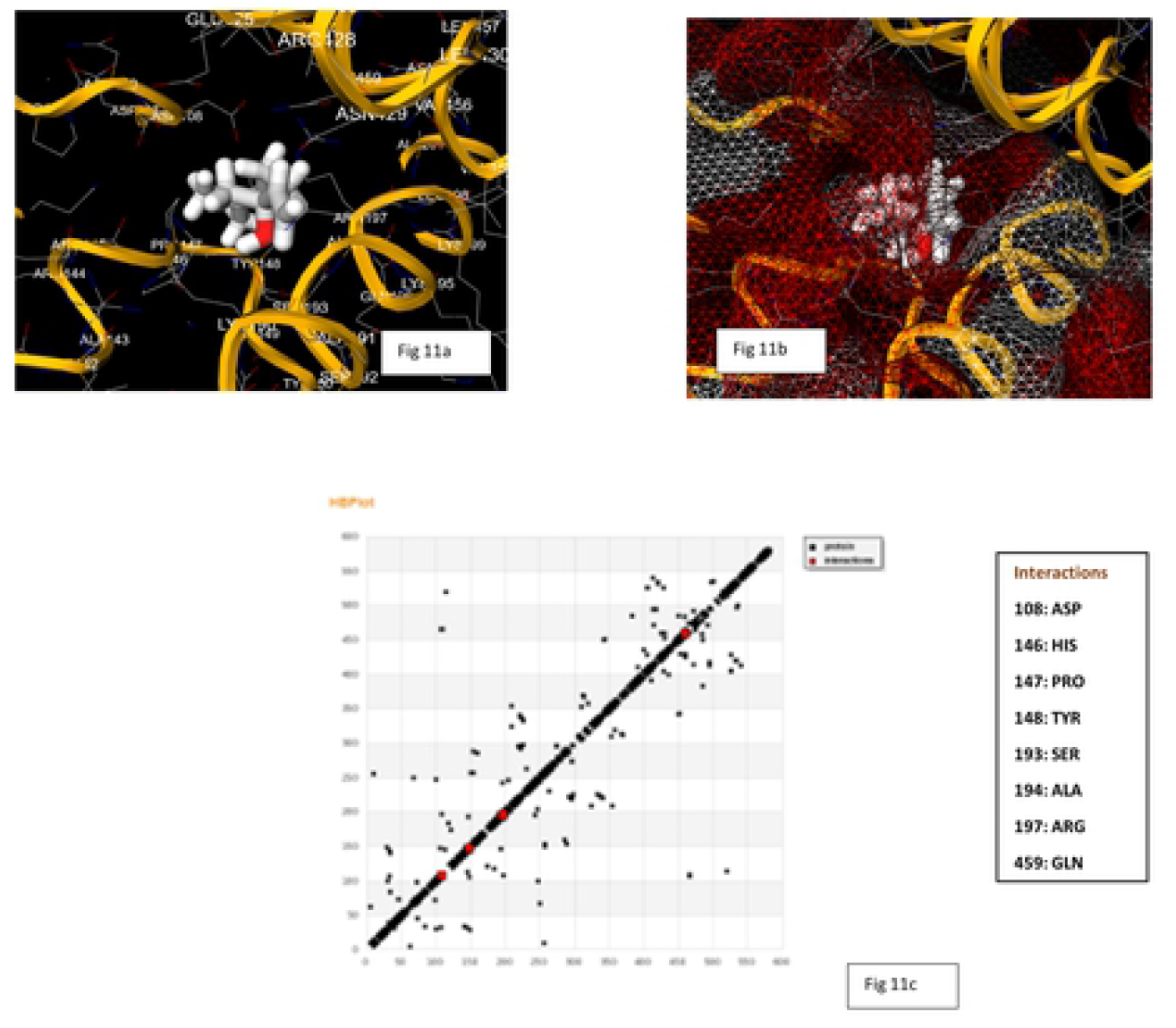
Fig 11(a). Docked orientation of linalool at the binding site of HSA with bonding residues showing stick model in yellow and ligand in white color, Fig 11(b). Binding mode of HSA with linalool binding to the HSA surface residues network like, Fig 11(c). HB Plot of linalool in the binding site of HSA.

The binding images of the predominate configuration of carvacrol–HSA complex is shown in Fig 11d and 11e, and the results of free binding energy and other values are given in Table 5. Moreover, as shown in Fig. 11d, the binding results indicate that carvacrol is very close to the binding site amino acid residues Asp108, His146, Pro147, Tyr148, Ser193, Ala194, Arg197, Gln459 at site I in subdomain IIA. The hydrogen bonding between carvacrol and HSA (Asp108) is also responsible for maintaining the stability of this complex. Whereas, the aromatic moiety of the carvacrol shown to engage the Tyr148, Ala194 and Pro147 for hydrophobic interaction. The other non-bonded interaction involves Arg197, Gln459, Ser193. The similar value of interaction has been also reported on close inspection of the HB plot of the carvacrol and HSA as shown in Fig 11f Carvacrol binds to the active site of HSA with minimum binding energy (ΔG) −7.1 kcal/mol (Table 5).

**Fig 11.**
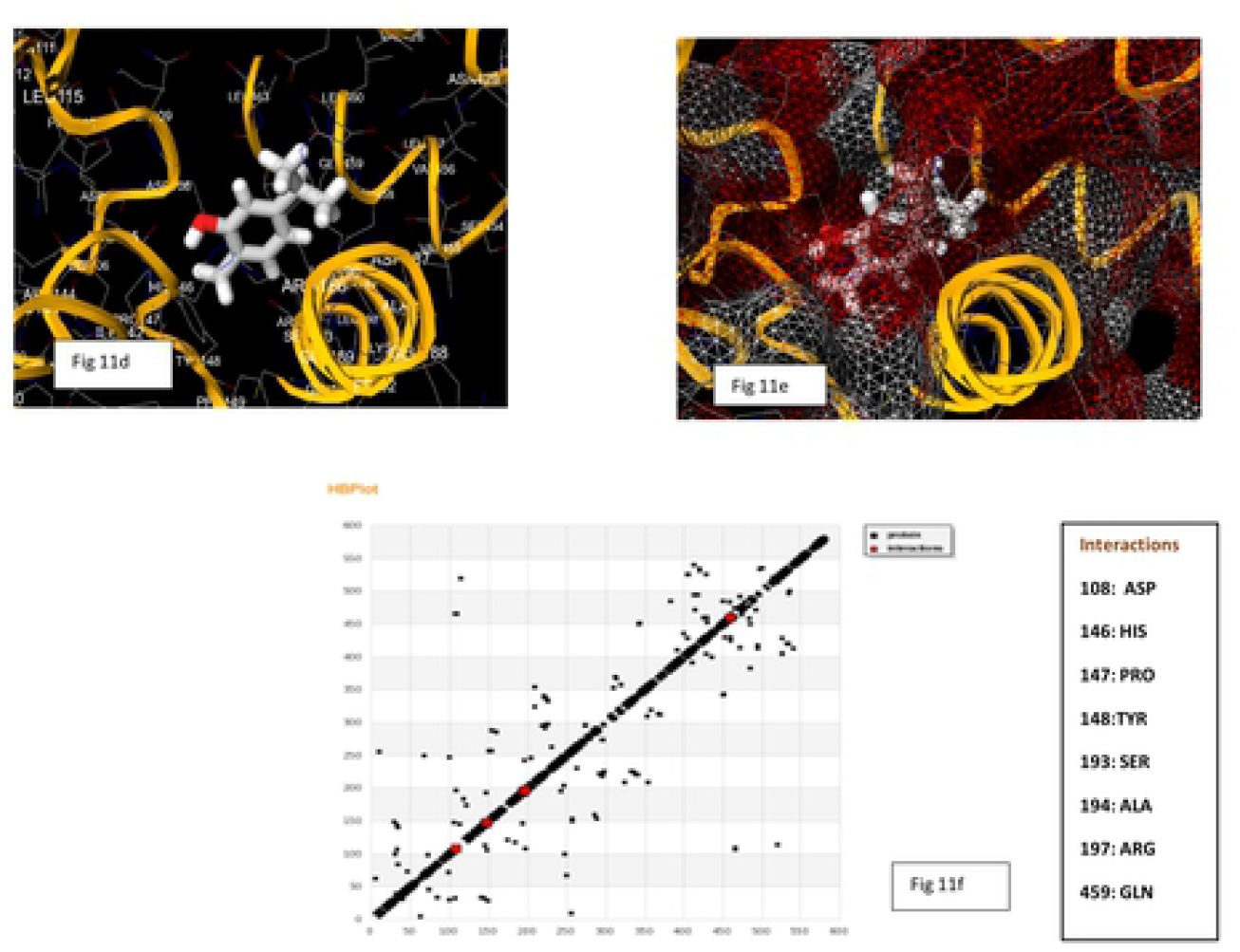
Fig 11(d). Docked orientation of linalool, carvacrol and thymol respectively at the binding site of HSA with bonding residues showing stick model in yellow and ligand in white color, Fig 11(e). Binding mode of HSA with linalool, carvacrol and thymol respectivey binding to the HSA surface residues network like, Fig 11(f). HB Plot of linalool, carvacrol and thymol respctively in the binding site of HSA.

The binding images of the predominate configuration of thymol–HSA complex is shown in Fig 11g and 11h, and the results of free binding energy and other values are given in Table 5. Moreover, as shown in Fig 11g, the binding results indicate that thymol is very close to the active site amino acid residues Asp108, His146, Pro147, Tyr148, Ser193, Arg197, Gln459, Val462, and Leu463 at site I in subdomain IIA. The hydrogen bonding between thymol and HSA (Ser193) is also responsible for maintaining the stability of this complex. Whereas, the aromatic moiety of the thymol shown to engage the Pro147, Val462, Leu463 for hydrophobic interaction and for cation-pi interaction His146. The other non-bonded interaction was also observed with the involvement of Arg197, Asp107, Gln459 and Tyr148. The similar value of interaction has been also reported on close inspection of the HB plot of the thymol and HSA as shown in Fig 11i. It was found to be nicely bounded into the active site of HSA with minimum binding energy (ΔG) −4.77 kcal/mol (Table 5).

**Fig 11.**
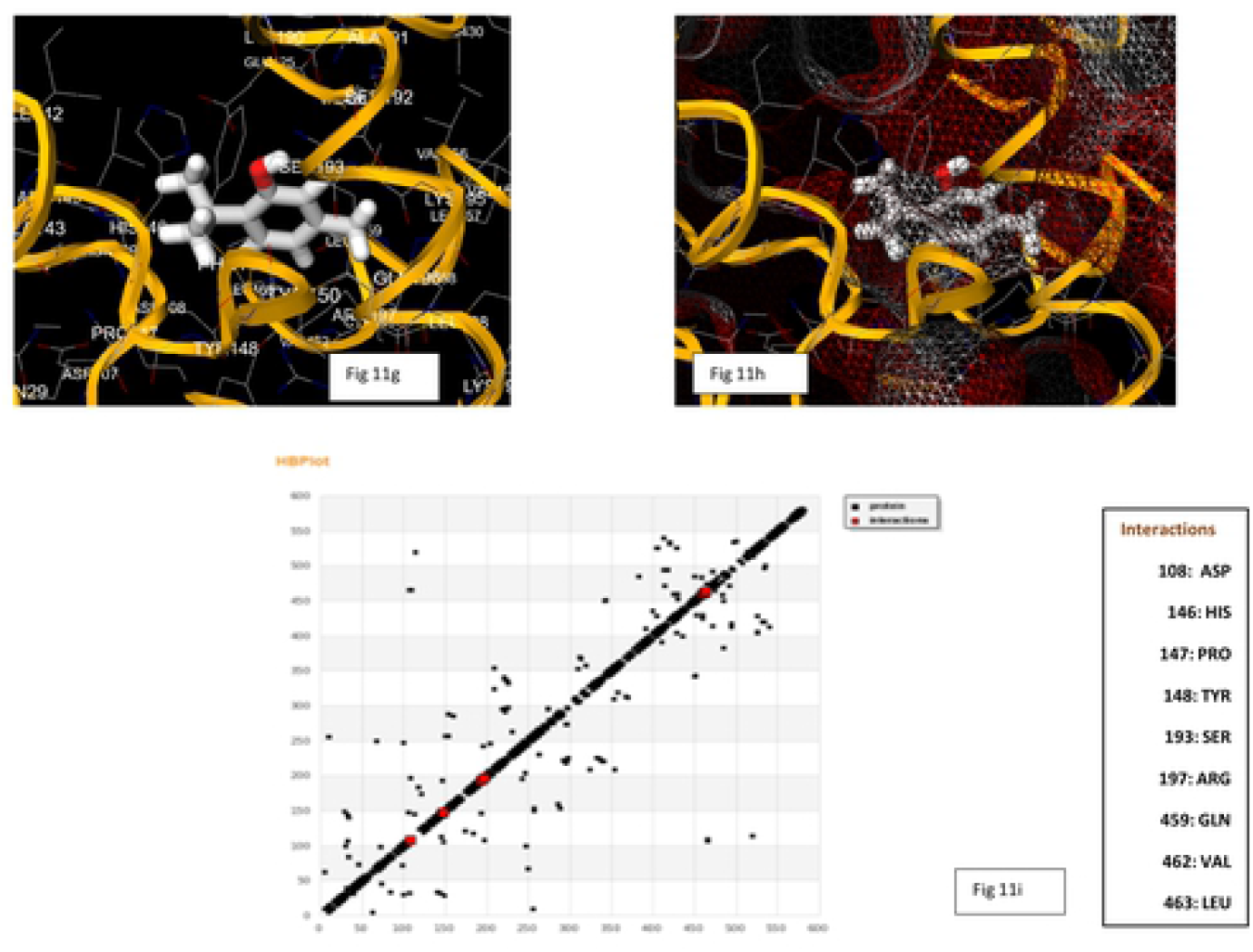
Fig 11(g). Docked orientation of thymol at the binding site of HSA with bonding residues showing stick model in yellow and ligand in white color, Fig 11(h). Binding mode of HSA with thymol binding to the HSA surface residues network like, Fig 11(i). HB Plot of thymol in the binding site of HSA.

Molecular docking studies suggested that the various van der Waals, covalent, Pi alkyl, carbon-hydrogen, and electrostatic interactions are the critical force for holding of all three tested ligands together with HSA. Therefore, it is clearly suggested that hydrogen bonds and hydrophobic forces were predominantly involved to stabilize the linalool–HSA, carvacrol–HSA and thymol–HSA complex as it had shown good binding energy for HSA.

## 4. Discussion

Compared to bacterial pathogens, fungi are evolutionary close to humans and thus have limited unique targets for drug discovery. Limited classes of antifungals currently available are sterol biosynthesis inhibitors (azoles and allylamine), cell wall inhibitors (polyenes and echinocandins) and DNA synthesis inhibitors (flucytosine). These existing antifungals are further besotted with problems of toxicity, undesirable drug interaction and emergence of drug resistance (26). Phytochemicals identified from traditional medicinal plants present opportunity for the development of novel therapeutic strategies. Recently, it has been shown that linalool exerts antifungal activity by disrupting membrane integrity, inhibiting germ tube formation, arresting the cell cycle of planktonic *C. albicans and* displaying activity against the cells of this yeast in biofilms (27). Lipase assays have revealed that carvacrol and thymol suppress enzymatic activity in *S. aureus* and inhibited production of staphylococcal enterotoxins. The reduction in the enzymatic activity (lipase) of the cells and in the synthesis of enterotoxins most likely occurred due to a prevention of protein secretion, which could have been a consequence of changes in the physical nature of the staphylococcal cytoplasmic membrane (28–30). Mostly all these processes are intimately associated with lipases. Such phenotypic modifications might possibly arise as a result of interactions between the phenolic compounds and enzyme. Studies have reported the chemical composition, phenolic content, antioxidant and anti-lipase activity of oregano essential oils against Candida Antarctica and Pseudomonas fluorescens with IC50 5.09 and 7.26 ugmL-^1^ respectively (31). The compounds found in oregano essential oil are thymol (8.00%), carvacrol (4%) and linalool (2.16%) etc. Its mechanism of action is variously reported as inhibition of ergosterol biosynthesis, disruption of membrane integrity, inhibition of yeast to hyphal transition and plasma membrane-ATPases activity. These processes are intimately associated with membrane and lipases.

Various virulence factors – lipase, proteinase, α amylase, phospholipase, pectinase, biofilm, hemolysin cause degradation of tissue carbohydrate (α–amylase), protein (proteinase), phospholipids (phospholipase), pectin (pectinase), and lipids (lipase). Lipases have been shown to influence growth, morphology, adherence and dissemination of fungal cells across host (32). Biofilm formation helps prevention of phagocytosis of pathogens and allows exponential growth. Hemolysin causes lysis of red blood cells.

Lipases in recent years have emerged as important pathogenic factors. Lipase activity is required for colonization and persistence of bacterial pathogens (33). Mycobacterium tuberculosis relies on lipases to hydrolyse host lipids as energy source (34). Fungal lipases thus can be explored as new therapeutic target. Fungal lipase share some structural and functional homology with human lipases but have low sequence homology and each lipase group possesses characteristic 3-D structure and catalytic properties (35). Fungal lipase as compared to mammalian lipase lack c-terminal domain and function as monomers (36). This may impart uniqueness and selectivity of target.

In this study we select three phytochemicals thymol, carvacrol and linalool which affect fungal growth and / or virulence and which are reported in literature to either directly or indirectly affect lipases or lipase related processes. Based on premise that a therapeutic molecule must disrupt structure of its target protein (i.e. lipase) but at the same time must have distribution in the system by binding to carrier proteins like HSA, interaction studies of all three phytochemicals have been systematically carried out on lipase, HSA.

Lipase showed approximately 50% activity at about 1.5mmoles/litre, of thymol, carvacrol and linalool respectively. However, for a natural molecule to be classified as therapeutic agent against fungal infection, it must show direct interaction to the target and its disruption. Accordingly, we perform binding studies of thymol, carvacrol and linalool with lipase. At the same time natural molecules must also bind to carrier molecule like HSA but should not disrupt their structures.

The UV-Vis spectroscopy gives evidence of the formation of lipase-thymol, lipase-carvacol and lipase-linalool, complex at the ground state. In case of HSA formation of the HSA -thymol, HSA – carvacrol and HSA – linalool complex is seen at the ground state.

Fluorescence spectroscopy gives insight into changes in tertiary structure of proteins following complex formation. Increase in fluorescence intensity attributed to decrease in rotational freedom in hydrophobic cavity occupied by aromatic ligand has been reported (37). Fluorescence spectra of lipase in presence of GdmHCl shows prominent red shift indicating denaturation. Similar results have been reported by other researchers for porcine and rice brown lipase at similar concentration (38). Thus, quenching binding constants of over 10^5^ M^-1^ are taken to be significantly strong (39, 40). Prominent red shift in on reset of lipase indicates major changes in tertiary structure and disruption of native structure. Strong binding of lipase to thymol is evident from *K_a_* of 2.6 x 10^9^ M^-1^as compared to its relatively weak binding to carvacrol (4.66 x 10^7^ M^-1^) and linalool (5.3 x 10^3^ M^-1^). Number of binding sites showing the stoichiometry of association process on lipase is found to be 2.52 (thymol) as compared to 2.04 (carvacrol) and 1.12 (linalool). These ‘n’ values obtained through double logarithmic plot of steady state fluorescence measurement, however, mean the stoichiometry of association process (41). These ‘n’ values suggest that stability of the lipase –thymol complex is significantly more as compared to lipase – carvacrol and lipase -linalool complexes which may have bearing on therapeutic selectivity of these ligands for lipase (42). Thus, the strength and stability of binding of three tested phytochemicals to lipase is in the order: Thymol > Carvacrol > linalool. Thymol and carvacrol can be classified as strong binders whereas linalool is a weak binder.

Further the secondary structure formation aimed by CD spectra results, following 24 hours incubation at 25°C, thymol, carvacrol and linalool revealed to cause decrease in negative ellipticity for lipase indicating loss in α helical structure as compared with the native protein. This indicates disruption of the hydrogen-bonding networks due to binding between secondary structural elements of the polypeptide chain of lipase and phytochemicals. Far - UV CD spectra of lipase in presence of 0.5M GdmCl, a known denaturant, also shows lowering in the negative ellipticity which comprise loss in secondary structure of lipase (38). Similar trends have been reported by other investigators for porcine and *Rhizopus nivens* lipase at comparable concentrations (43).

The lowering in negative ellipticity following binding of phytochemicals is observed in the order: Thymol > Carvacrol > Linalool.

Results of fluorescence and CD spectroscopy taken together would suggest thymol and carvacrol to be more profound disrupter of lipase structure at comparable concentrations. Whereas linalool is poor disrupter of structure.

Fluorescence spectra of binding of all phytochemicals: thymol, carvacrol and linalool with Human Serum Albumin (HSA) caused blue shift which suggests the compaction of the HSA structure. Association constant of thymol with HSA is 9.6 x 10^8^ M^-1^ and along with ‘n’ value of 2.41 suggests strong association and stable complex formation. Observed blue shift would however suggest in change in tertiary structure towards compaction. Linalool and carvacrol binding association values in the range of 10^4^ M^-1^ indicate somewhat poor binding and ‘n’ values of 1.43 and 1.29 somewhat less stable. Linalool-HSA and carvacrol-HSA complexes It may be noted that fluorescence spectra HSA in presence of GdmCl cause red shift with HSA (44).

CD spectroscopy of HSA with thymol, carvacrol and linalool shows increase in negative ellipticity for HSA suggesting gain in alpha - helical structure. Carvacrol shows the least value of MRE as compared to other two molecules indicating highest gain of α-helical structure among the all molecules. Next thymol and linalool show almost equal gain of α-helical structure. Fluorescence spectroscopy results take a along with CD spectroscopy would suggest thymol as good and stable binder of HSA. In addition, these molecules do not disrupt the structures of HSA making the ligand suitable for dispersal in blood milieu.

These results taken together with lipase binding would suggest thymol and carvacrol as good binder and disrupter of lipase structure along with reasonable binding with HSA for distribution. Interestingly these molecules do not disrupt structure of HSA and have only moderate binding which may make the association reversible. This fact is of vital importance for potential therapeutic agents.

And finally docking results also give a clear insight suggesting strong binding of thymol, carvacrol and linalool with lipase having free energy of binding −7.1 kcal/mol, −5.0 kcal/mol and 5.28 kcal/mol respectively. Thymol, carvacrol and linalool also bind to HSA with good affinity showing −4.77 kcal/mol, −4.77 kcal/mol and −4.21 kcal/mol of free energy of binding respectively.

## 5. Conclusion

Among the fungal virulence factors, extracellular lipase can be a very promising candidate for combating fungal diseases. In this study fungal *Aspergillus niger* lipase is explored as target of thymol, carvacrol and linalool using UV-visible, fluorescence and CD spectroscopy. Binding studies of the phytochemicals have also been carried out on Human Serum Albumin to analyse their therapeutic potential. All three phytochemicals caused 50% inhibition of lipase at around 1.5mM. UV-vis spectra suggested that all phytochemicals formed stable complexes with lipase and HSA. Sequence of binding affinity with *Aspergillus niger* lipase was thymol (10^9^) > carvacrol (10^7^) > linalool (10^3^). Increase of Mean residue ellipticity (MRE) also followed the same order suggesting that thymol and carvacrol are best binders and disrupters of lipase structure. Sequence of binding affinity with HSA was thymol (10^8^) > linalool =carvacrol (10^4^), decrease of MRE (more –ve) was observed in the order: carvacrol> thymol > linalool. Thus, best binder is thymol whereas all three are good compacters of HSA structure.

Best anti-lipase molecule for human system identified on basis of lipase structure disruption and HSA structure conservation is thus thymol. Docking studies also indicate strong binding of thymol and lipase with binding energy of −7.1 kcal/mol.

## Acknowledgement

FN gratefully acknowledges financial support in form of Maulana Azad Minority Fellowship from University Grants Commission, India.

## Supporting information

**S1(A-C) Figs. Stern–Volmer plot (lipase–linalool, lipase–carvacrol and lipase–thymol).**

**S2(A-C) Figs. Double-logarithmic plot for the fluorescence intensity of lipase with various concentrations of the linalool, carvacrol and thymol.**

**S3(A-C) Figs. Stern–Volmer plot (HSA-linalool, HSA-carvacrol and HSA-thymol).**

**S4(A-C) Figs. Double-logarithmic plot for the fluorescence intensity of HSA with various concentrations of the linalool, carvacrol and thymol.**

**S5 Fig. Predicted structure of Lipase *Aspergillus niger* (*A niger* lipase).**

